# A possible molecular mechanism for mechanotransduction at cellular focal adhesion complexes

**DOI:** 10.1101/2020.12.16.423152

**Authors:** Jichul Kim

## Abstract

Mechanotransduction at focal adhesion complexes is key for various cellular events. Theoretical analyses were performed to predict a potential role of lipid membranes in modulating mechanotransduction at focal adhesions. Calculations suggest that the nanoscale geometric changes and mechanical pulling applied on lipid membranes affect the generation of cellular traction forces and signaling transduction at focal adhesions. This work provides predictions on how lipid membranes contribute to mechanotransduction at cellular focal adhesions.

**Significance statement:** Focal adhesion machinery formed across cell membranes orchestrates a variety of signaling and adhesive molecules to function for important cellular physiologies. Although there are evidences that lipid membranes are involved in mechanical transduction at focal adhesions, how the detailed mechanical response of membranes contributes to the process is not identified yet. With numerous data previously identified, predictions made by theoretical modeling suggest that the nonlinear pulling response of lipid membranes serves as a key factor to interpret mechanotransduction at focal adhesions.

## Introduction

Cells interact with their environments and generate traction forces through their adhesive molecular machinery formed across cell membranes (1). This collective molecular complex, referred to as the focal adhesion (FA), is sequentially composed of extracellular matrix proteins, adhesive integrin receptors inserted in membranes, adaptors such as talins and vinculins in the cytoplasm, and stretched by the contraction of actomyosin motors in the cytoskeleton. Forces transmitted through these molecules generate biological signals that affect cellular growth, differentiation, migration and tumor metastasis (2). Therefore, understanding FAs is central for various physiological and pathological processes.

Recent mechanical measurements on FAs identified their underlying molecular and biophysical mechanism from at the cellular level to at the single-protein level, and thus provided a multiscale perspective for the mechanism of FAs (3-10). Nevertheless, an integrated interpretation on how the individual molecular response affects the generation of the complex cellular mechanical behavior and the signaling transduction at FAs is still elusive. Moreover, the effect of the mechanical stretching of lipid membranes, a potentially important component at FAs, is not fully studied yet.

In this work, the combined nanomechanical response of lipid membranes and talin proteins was investigated. By connecting the membrane-talin response to other FA mechanisms, cellular traction forces on elastic substrates were also calculated. Furthermore, vinculin binding to force-stretched talins and its relation to the nanoscale mechanics of the membrane-talin complex were investigated. Overall, this work provides an idea for the integrated FA molecular machinery, by emphasizing a crucial role of lipid membranes in modulating mechanotransduction.

## Methods

### Modeling the mechanical response of lipid membranes at the membrane-talin complex

Among various molecules, two key elastic components at FAs were assumed in this model; lipid membranes and talins (1). The membrane is linked to the cytoskeleton through integrin-bound talin molecules, and they share the mechanical extension *E_mem-talin_* (Fig. 1A and B). The lipid membrane where clustered integrins are inserted, was described as coarse-grained continuum by using a finite element method (FEM) introduced in a recent investigation (11).

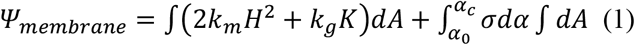

**Figure 1.**
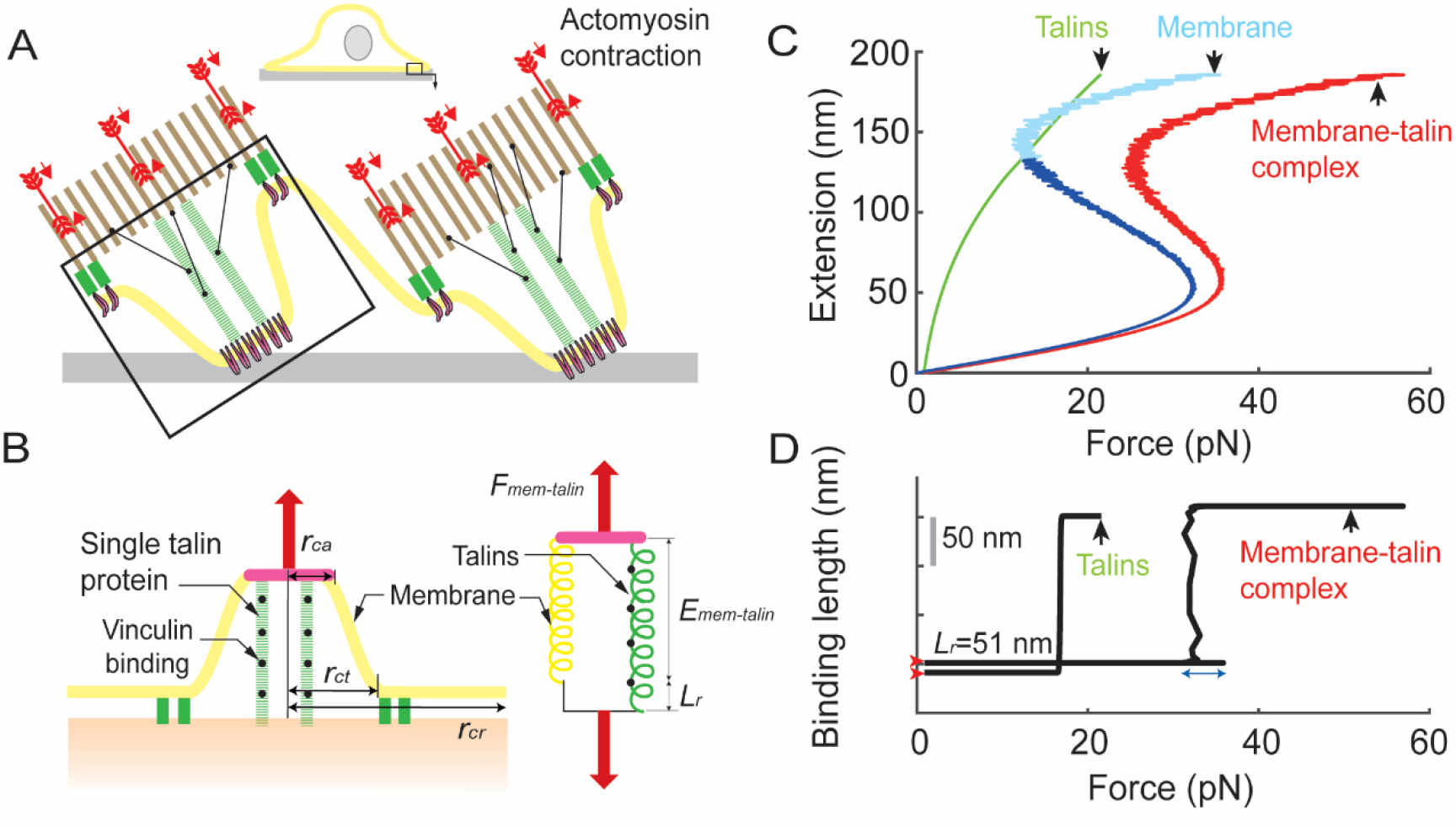
Extension vs. force and vinculin binding length vs. force curves of the single membrane-talin complex. (A and B) Illustration for a unitary adhesion complex composed of clustered integrins (magenta), lipid membranes (yellow), and talin proteins (green). Forces applied on integrins and actomyosins (brown+red) stretch the membrane-talin complex. The extension results in the binding of vinculin molecules (black) to talins. (C) The extension vs. force calculation for the membrane-talin complex (red). The membrane component (cyan) and the component from talins (green) were also plotted separately. The curve in red is the sum of the two curves in cyan and green. *r_ca_* = 57 nm, *r_ct_* = 158 nm and *r_cr_* = 1750 nm were used, and resulting *N_talin_* is approximately *N_talin_* ≈ 9. The initial talin force is not zero due to *L_r_*. The membrane response in a smaller extension regime is used for the calculation in Fig. 3C middle panel (blue). (D) The vinculin binding length vs. force calculation for the membrane-talin complex and for the case without the mechanical rigidity of membranes (i.e., Talins). The resting binding length *L_r_* is 51 nm (red arrows). Δ*G*=252.9*k_b_T* and Δ*G*= 1069*k_b_T* for Talins and Membrane-talin complex, respectively.

**Figure 2.**
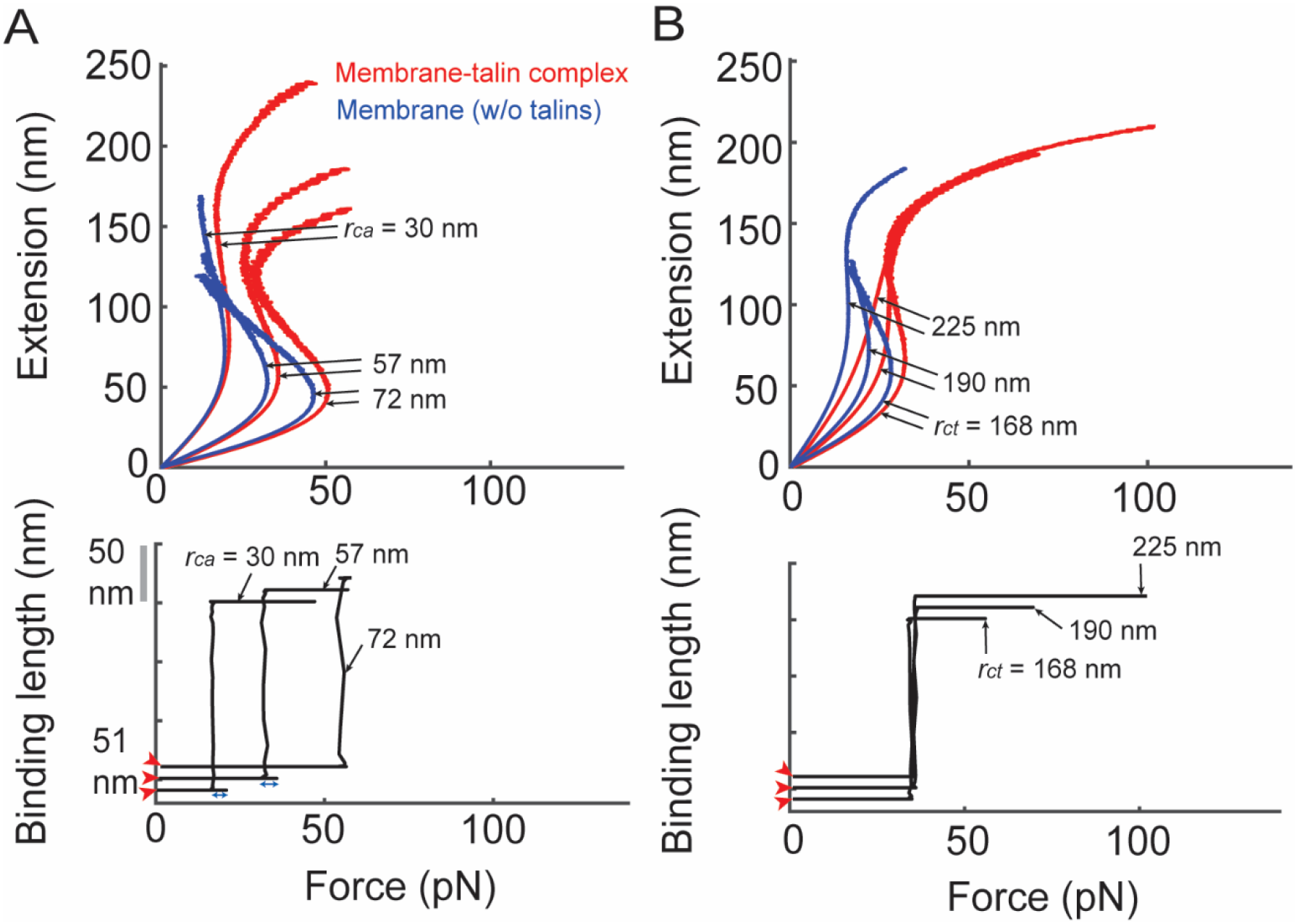
Modulation of extension vs. force and vinculin binding length vs. force curves for the single membrane-talin complex. (A) Mechanical responses of the membrane-talin complex (red) and the complex without talin molecules (blue) by varying the *r_ca_* value. *r_ct_* = 158 nm was used for all six calculations. The vinculin binding length curves were plotted for the membrane-talin complex (bottom panel). The resting binding length *L_r_* is 51 nm (red arrows). Δ*G* = 638.4*k_b_T* and *N_talin_* ≈ 3 (*r_ca_* = 30 nm); Δ*G* = 1069*k_b_T* and *N_talin_* ≈ 9 (*r_ca_* = 57 nm); and Δ*G* = 1445.2*k_b_T* and *N_talin_* ≈ 15 (*r_ca_* = 72 nm). (B) Responses with different *r_ct_* values. *r_ca_* = 57 nm for all six calculations. Δ*G* = 1021.9*k_b_T* and *N_talin_* ≈ 9 (*r_ct_* = 168 nm); Δ*G* = 912*k_b_T* and *N_talin_* ≈ 9 (*r_ct_* = 190 nm); and Δ*G* = 759.7*k_b_T* and *N_talin_* ≈ 9 (*r_ct_* = 225 nm). See Fig. S3 for additional sensitivity analyses.

In equation (1), the energy functional *ψ_membrane_* is expressed with energies associated with the mean *H* and Gaussian *K* curvatures of the projected area of the membrane *A* (12, 13). The mean curvature at certain points of the area is 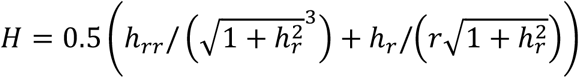 where *h* is the height function of the projected membrane shape, and *r* is the rotational-symmetric radial function. The first and second derivatives of *h* with respect to *r*, *h_r_* and *h_rr_*, were expressed with respect to the parametric coordinate *s*[0,1] in the framework of parametric FEMs by using *h_r_* = *h_s_*/*r_s_* and *h_rr_* = *h_ss_*/*r_s_*^2^ − *h_s_r_ss_*/*r_s_*^3^, respectively (11). *k_m_* and *k_g_* are the bending modulus and the Gaussian curvature modulus, respectively. In performing the variational calculation, the Gaussian curvature energy term in equation (1) was omitted, based on the Gauss-Bonnet theorem (14). For simplicity, *k_m_* was assumed to be constant in this work (15–18).

The functional *ψ_membrane_* contains an area strain energy term. The strain energy density 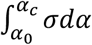 in equation (1) can be calculated by integrating the surface tension *σ* vs. area strain *α* relation from the resting reference strain *α*_0_ to the strain under consideration *α_c_*. Two smooth expressions for the surface tension are *σ* = *σ*_0_ *exp*(8*πk_m_α*/(*k_b_T*)) for *α* ≤ *α_cross_* and *σ* = *K_app_*(*α* − *α_cut_*) for *α* > *α_cross_*, where *α_cut_* and *α_cross_* are cut-off and cross-over strains, respectively. *K_app_* is apparent area stretching modulus (11, 19, 20). The strain is 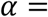 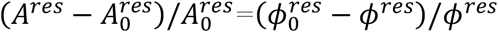 where *A^res^* is the projected area of the membrane reservoir determined by *r_cr_* − *r_ct_* and *s*[0,1]; 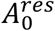 is the projected area of the lipid reservoir at the resting reference configuration; *ϕ^res^* is the uniform lipid number density; and 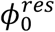 is *ϕ^res^* at the resting reference configuration. *k_b_* is the Boltzmann constant and *T* is the temperature in Kelvin.

The boundary conditions are (*r*(0), *h*(0)) = (*r_ca_*, *E_mem-talin_*), (*r_s_*(0), *h_s_*(0)) = (*r_ct_* - *r_ca_*, 0), (*r*(1), *h*(1)) = (*r_ct_*, 0), and (*r_s_*(1), *h_s_*(1)) = (*r_ct_* - *r_ca_*, 0). *r_ca_* defines the radius of a rigid area where clustered integrins are inserted (10). *r_ct_* defines the size of tented membranes in response to forces on the stiff area. Image data showing membrane curvatures at FAs may support this parameterization (21, 22). *r_cr_* was introduced to assume the radial area for the lipid reservoir. *r_cr_* may not represent a physical geometry, but it simply defines the number of lipids available in mechanical stretching (Fig. 1A and B). The curvature change was not considered for the region assumed by *r_cr_* (11).

Calculations for the mechanical response i.e., the extension *E_mem-talin_* vs. force *F_mem_* response, were performed by directly employing the finite element method provided in the previous work (11). In short, the function *h*(*s*) and *r*(*s*) in the weak form of equation (1) were parameterized with the B-spline function, and the resulting nonlinear equation system was solved by using the Newton-Raphson method. The stationary of equation (1) was assumed when the Euclidean norm of the difference of two subsequent solution vectors converged to a certain value. In increasing *E_mem-talin_*, an estimate for the lipid number density for the *k*+1^th^ step was calculated from the *k*^th^ step, when the calculated projection area of the tented membrane in the *k*^th^ step is greater than that in the *k*-1^th^ step. Here, *k* is the index for the step extension. (See ref. (11) for the details of the finite element method).

### Modeling vinculin binding to talin at the membrane-talin complex

Talin proteins are known to interact with integrins and actins at FAs (23, 24) (Fig. 1A and B). The total number of talin rods connected to integrins in the single membrane-talin complex *N_talin_* can be *N_talin_* = *C_talin_N_integrin_*. The number of integrins *N_integrin_* in the area defined by *r_ca_* can be estimated by assuming the area occupied by a single integrin pair with the radius *r_integrin_* i.e., *N_integrin_* = (*r_ca_*/*r_integrin_*)^2^ (25, 26). *C_talin_* is the fraction of talin molecules with respect to the integrins in *r_ca_*. The mechanical extension vs. force *F_talin_* response of single full length talins was assumed by smoothly extrapolating a previous experimental data measured on a larger extension regime (9) (Fig. S1). Finally, forces applied on the single membrane-talin complex *F_mem-talin_* can be *F_mem-talin_* =*F_mem_*+*N_talin_F_talin_* (Fig. 1B red arrows).

Talin rods convert mechanical inputs into biological protein interactions. Vinculin binding to force-stretched talins is a major mechanical transduction event at FAs (27). A statistical description was introduced to investigate the vinculin binding event with respect to the continuous mechanical pulling of the membrane-talin complex, and therefore to plot the length of vinculin binding to talins. The total potential energy of the membrane-talin complex *ψ_total_* was modeled by assuming two-state variable *λ* as in equation (2).

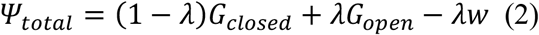

Here, *λ* = 0 and *λ* = 1 represent the closed and open state of the vinculin binding sites in talins, respectively. The two-state condition might not be enough, as single talin rods have multiple vinculin binding sites connected in series with potentially different open and closed configurations (28). However, the model provided a simple framework to evaluate the mechanical transduction at the membrane-talin complex with the two-state assumption. *G_closed_* and *G_open_* are the internal energies of the single membrane-talin complex in closed and open configurations of the vinculin binding sites, respectively. The value *w* is the external work, and it was calculated by integrating the force vs. extension response of the membrane-talin complex. By substituting the total energy in equation (2) into the Boltzmann function, the probability of finding the open state of talins *p_λ_*=1 can be obtained from equation (3) where *β* = 1/(*k_b_T*) (29).

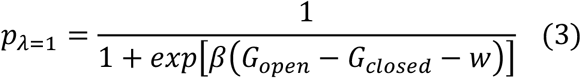

The parameter value Δ*G* = *G_open_* - *G_closed_* was obtained by integrating the force vs. extension curve of the single membrane-talin complex from a resting extension to a reference extension for talin activation. The reference extension length *L_ref_* = 160 nm was used (5). With different *r_ca_*, *r_ct_* and *r_cr_* values, Δ*G* was differently calculated since the membrane energy and the number of talin proteins are different for the single membrane-talin complex. With *r_ca_* = 57 nm, *r_ct_* = 158 nm and *r_cr_* = 1750 nm, Δ*G* was 1069*k_b_T* where 252.9*k_b_T* and 816.1*k_b_T* were generated from nine talins and membranes within the membrane-talin complex, respectively. This Δ*G* value demonstrated that the membrane mechanical response is an important factor in this model for mechanotransduction at FAs. Finally, the model assumed that the length of vinculin binding to talins *L_binding_* is proportional to the probability of finding the open state of the vinculin binding sites *p_λ_*=1 i.e., *L_binding_* = *L_r_* + *p*_*λ*=1_*L_ref_*. The constant *L_r_* = 51 nm is the vinculin binding length in the resting configuration, or it can simply define the resting length of talins (5, 30) (Fig. 1B). The elasticity of vinculin molecules was not considered in this work.

### Modeling cellular traction forces from the mechanical response of the membrane-talin complex

To investigate how the membrane-talin response contributes to the generation of cellular traction forces, the stiffness of extracellular substrates was modeled as in equation (4) by directly following previous investigations (8, 31) (Fig. 3A and B).

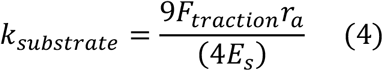

**Figure 3.**
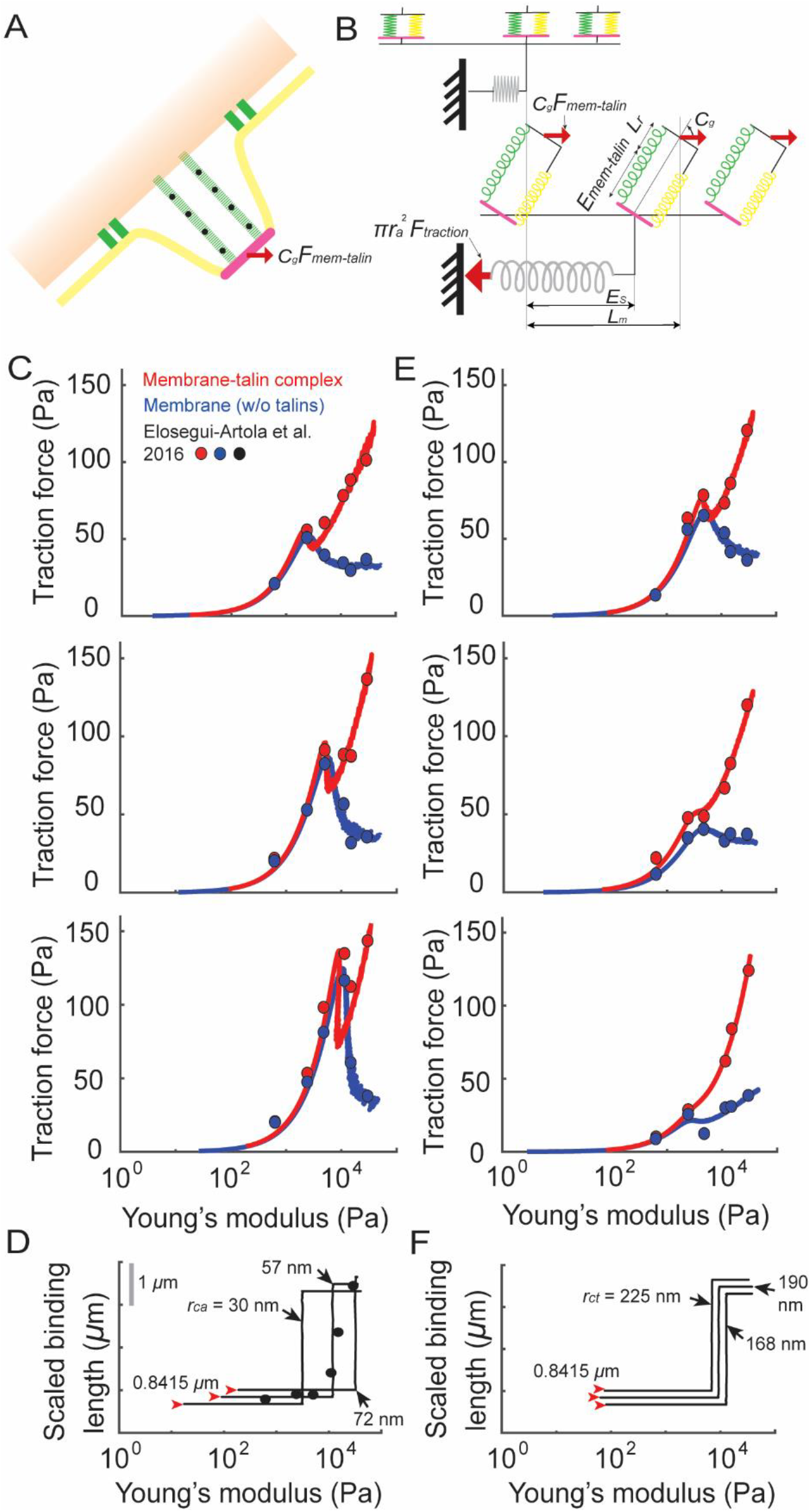
Cellular traction force vs. substrate stifness curves calculated from the responses of the membrane-talin complex. (A and B) Illustration for how the single membrane-talin complex is linked to the generation of cellular traction forces at focal adhesions. (C) Traction forces calculated by using data in Fig. 2A top panel. The curves were matched to experimental measurements obtained by treating the different amounts of fibronectin on elastic substrates. (D) The responses for the vinculin binding length scaled from the data in Fig. 2A bottom panel. The calculation with *r_ca_* = 57 nm was directly matched to the experimental data. The scaled resting length was 0.8415 μm (red arrows). (E) Traction forces calculated by using data in Fig. 2B top panel. The curves were matched to measurements obtained with the treatment of the different amounts of the integrin-binding peptide GPen. (F) The responses for the vinculin binding length scaled from the data in Fig. 2B bottom panel. Red: responses from the membrane-talin complex. Blue: responses without talins. Circles: measurements from ref. (8).

Here, *k_substrate_* is the Young’s modulus of substrates. *F_traction_* is the traction force applied on an area with the radius *r_a_*, and *F_traction_* = *F_mem-talin_N_mem-talin_C_g_*/(*πr_a_*^2^) where *N_mem-talin_* = *r_a_*_2_/*rct*^2^ (or *N_mem-talin_* = 1 when *r_a_* = *r_ca_*) is the number of the membrane-talin complex in the continuous adhesion area. *C_g_* is the geometric coefficient to account for the tile of the molecular complex with respect to substrates (see Fig. 3B). *E_s_* is the lateral extension of substrates and *E_s_* = *L_m_*-*C_g_*(*E_mem-talin_*+*L_r_*)>0. Here, *L_m_* is the size of serial stretching of substrates and the membrane-talin complex (with talin’s resting length *L_r_*) in the surface-horizontal direction. Since the stretching of the FA molecular machinery results from the contraction of actomyosin motors (32), *L_m_*/*C_g_* may represent the size of the actomyosin contraction in the cytoskeleton. The minimum *L_m_*/*C_g_* value of 68.8 nm used in this study (for calculations in Fig. S4C) was greater than the maximum step size of 11 nm measured from single actomyosin motors (33). Finally, a free parameter *C_scaled-binding_*, that may suggest the alignment of multiple membrane-talin complexes at the bottom of cells, was introduced to calculate the vinculin binding length observed at cellular FAs by using *L_scaled-binding_* = *C_scaled-binding_L_binding_*. The parameter values used in this study are summarized in Fig. S5 and Table 1. Calculations and analyses were performed by using MATLAB (MathWorks^®^).

**Table 1.**
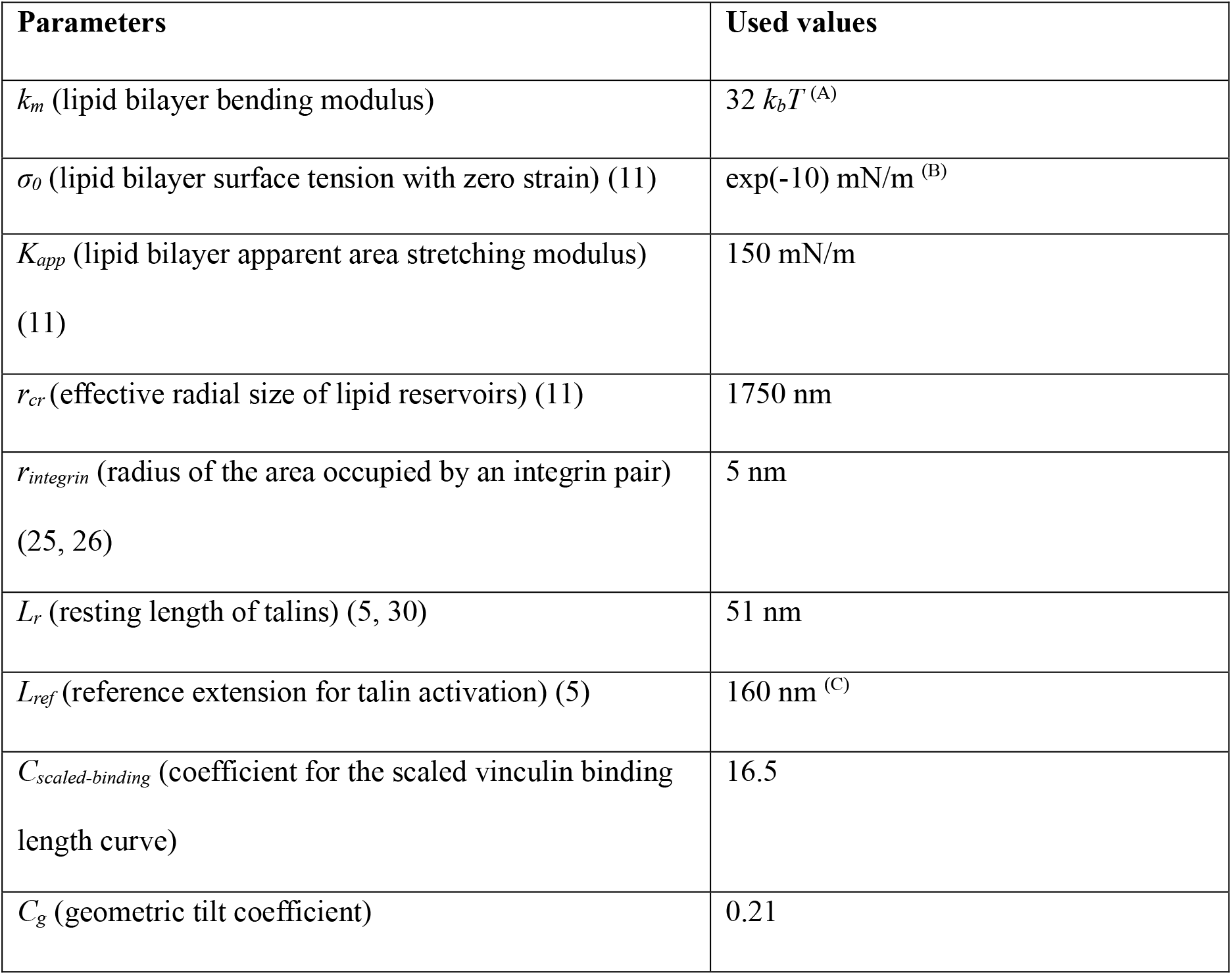
Summary of parameter values. (A and B) Sensitivity of *k_m_* and *σ_0_* on the extension vs. force response of the membrane-talin complex is supplemented in Fig. S2. (C) Resting length *L_r_* is not included in this value. See Fig. S5 for the use of other parameters

## Results and Discussion

### Nonlinear mechanical response of lipid membranes modulates vinculin binding to talins at the membrane-talin complex

The extension of the membrane-talin complex that resulted from changing force is shown in Fig. 1C. In this calculation, *r_ca_* = 57 nm was used (10), where approximately nine talin rods were connected in parallel with the membrane; *r_ct_* = 158 nm and *r_cr_* = 1750 nm were used (11). The response of the membrane-talin complex was nonlinear with the intermediate negative extension/force slope, and that was generated from the response of the membrane component. Forces applied on the complex were mostly conveyed to the membrane until the force became ~35.9 pN, and the extension of the membrane-talin complex was less than 60 nm in this region. Around 35.9 pN force, the calculation suggested an extension jump of the membrane-talin complex. Here, about 165 nm extension of talin rods within the complex is similar to a previous experimental observation (5).

The activation of mechanotransduction is plotted in Fig. 1D by calculating the vinculin binding length to talins, with forces applied on the membrane-talin complex. Without considering the mechanical rigidity of lipid membranes, the steep region of the vinculin binding length curve was located at around 16.8 pN (Fig. 1D, Talins). However, the curve was shifted to the higher force regime by considering the membrane connected in parallel with the talin rods (Fig. 1D, Membrane-talin complex). This result suggests that the deformation of membranes can modulate vinculin binding to talin rods when the FA molecular machinery is mechanically stretched. Interestingly, the steep region of the binding length curve and the region where the different vinculin binding lengths were predicted with one force value (Fig. 1D blue arrow around 31–35.9 pN) overlapped with the negative stiffness region (or the region for the extension jump) of the extension vs. force curve. Here the values 31–35.9 pN were similar to a previous measurement for the activation force of integrin signaling.

### Nanoscale geometric changes of lipid membranes modulate vinculin binding to talins at the membrane-talin complex

The idea that pulling of cell surface receptors can generate nonlinear nanomechanical responses was recently evidenced experimentally (11). Analyses suggested that the responses can be modulated by rigid components interacting with lipid bilayers (11). Therefore, the size of adhesion clusters and the level of the membrane-cytoskeleton interaction can serve important roles in modulating the mechanical response and the signaling transduction at FAs. To test these possibilities, *r_ca_* and *r_ct_* values were systematically varied in calculating the extension vs. force and vinculin binding length vs. force curves of the membrane-talin complex. As shown with red curves in Fig. 2A, increasing *r_ca_* resulted in responses that showed more predominant sigmoidal nonlinearity. The initial force peak i.e., the signature for the generation of the negative stiffness, was shifted to the higher force regime by increasing *r_ca_*. Shifting of the vinculin binding length curve was also identified, and the force peak of each extension vs. force curve approximately overlapped with the steep region of the binding length curve. In Fig. 2B, the nonlinear shape of the extension vs. force curve was also changed by varying *r_ct_* from 168 nm to 190 nm and 225 nm. It was demonstrated here that the shifting of the vinculin binding length curve in varying *r_ct_* with a given *r_ca_* of 57 nm was smaller than that resulting from varying *r_ca_* from 30 nm to 57 nm and 72 nm with a fixed *r_ct_* of 158 nm. In Fig. 2, responses of the unit complex without the talin molecules were also plotted (blue curves). Additional sensitivity analyses are shown in Fig. S3.

### The nonlinear mechanical responses and the nanoscale geometry changes of the membrane-talin complex affect the generation of cellular traction forces at FAs

By using the responses shown in Fig. 1C, traction force vs. substrate stiffness responses were generated in Fig. 3C middle panel. The responses demonstrated that the traction forces, calculated from the membrane-talin and membrane-without-talin responses, showed the similar nonlinear characteristics. It was remarkable that direct comparison between the calculations and previously measured data from living cells showed good agreements for both cases, with and without talins (8) (Fig. 3C, middle panel). The function of talin rods is closely related to the adhesive area and the Myosin II activity at FAs (see ref. (34)). Therefore, *r_a_* and *L_m_* were differently assigned for the traction force calculation of the membrane-talin and membrane-without-talin components (Fig. S5). The vinculin binding length curve for the membrane-talin complex in Fig. 1D was also compared to measurements in Fig. 3D (*r_ca_* = 57 nm and black dots). The result demonstrated that the Boltzmann function widely used in interpreting biological signaling systems (35, 36) can be also used for mechanotransduction at FAs. Similarly, as shown in Fig. 1C and D, the negative slope region of the force vs. stiffness response (Fig. 3C middle panel, red) overlapped with the steep region of the scaled response for the vinculin binding length (Fig. 3D, *r_ca_* = 57 nm).

The effect of *r_ca_* on cellular traction forces was then investigated. For this purpose, the membrane-talin and membrane-without-talin calculations using *r_ca_* = 30 nm and *r_ca_* = 72 nm shown in Fig. 2A were translated to plot the traction force vs. substrate stiffness responses in Fig. 3C top and bottom, respectively. Here different *r_a_* values were used, based on a recent identification that the increase of the size of integrin clusters is positively correlated with that of the total FA area (10). The *L_m_* was also varied since the FA area and the Myosin activity are closely interrelated (see ref. (34)) (Fig. S5). Gradual changes in the traction force responses were identified by increasing *r_ca_*. These changes in the traction forces showed good agreements with experimental measurements obtained by varying surface fibronectin density (8) (Fig. 3C), suggesting a tendency that the increase of fibronectin density can result in the increase of the integrin cluster size. In Fig. S6, fibronectin density vs. *r_ca_* relation was plotted. The shifting of the vinculin binding length curve to the high stiffness regime was identified in increasing *r_ca_* (Fig. 3D).

Figure 2B demonstrated that, with a fixed *r_ca_*, the initial stiffness of the membrane-talin response can be diminished by increasing *r_ct_*. This result provided a membrane-based hypothesis for a widely recognized pharmacological method using peptide sequences to modulate the integrin-ligand interaction (4, 37, 38). According to previous investigations, the peptide treatment to inhibit ligand binding to αvβ3, but not to α5β1 integrins, results in the inhibition of the stiffening response of normal cells similar to that observed in talin-depleted cells (4) as well as the change of the nonlinear force vs. stiffness response shape generated from talin-1-null cells (8). These results guided to investigate the relation between the peptide treatment and the effect of changing *r_ct_* in the mechanical response of the membrane-talin complex. To compare the calculations in Fig. 2B to traction force vs. substrate stiffness measurements from peptide-treated talin-1-null cells (8), the *r_a_* and *L_m_* values were systematically increased with the change of *r_ct_* (Fig. S5). Direct comparison between the calculations and the measurements showed good agreements (Fig. 3E). The result suggested that the integrin-binding peptide can modulate the size of the tented lipid membrane without directly affecting the size of integrin clusters. The explicit relation between the peptide concentration and *r_ct_* is shown in Fig. S6. Overall, it might be important to investigate whether and how the complex of α5β1 integrins and talin 2 molecules is stretched and slide on lipid membranes to modulate *r_ct_* when the FA is mechanically perturbed (39).

## Conclusion

Mechanical force generation and signaling transduction at focal adhesions (FAs) are key for various cellular physiologies. Several theoretical models were proposed to explain observed mechanical responses at FAs, based on dynamic bonding between different molecular components (7, 8, 40), as well as the unfolding property of adaptor proteins (8). In this work, another possibility is proposed by asking how lipid membrane mechanics contributes to mechanotransduction at FAs. The combined mechanical response of membranes and talins was nonlinear. According to sensitivity analyses, this nonlinearity can be modified with the nanomechanical geometry changes applied to the lipid membrane component, and that is linked to the vinculin binding activity to force-stretched talin rods. The results suggested that the membrane pulling mechanics determined by *r_ca_* and *r_ct_* can serve as an important factor to modulate mechanotransduction at FAs.

Cells are complex and involved in many active mechanisms related to each other. Therefore, the simple membrane parameters were insufficient; the model required several other parameters to translate the membrane-talin responses into the cellular traction forces. Nevertheless, the fact that these parameters were based on observations, as well as varied in a systematical way, may support their validity (Fig. S5). To demonstrate the importance of other parameters, analyses performed by varying *L_m_*, *C_talin_* and *r_a_*, with fixed membrane parameters, were supplemented in Fig. S4. It might be possible that other nanomechanical characteristics of membranes generated by the effect of multiple *r_cr_* (11), osmosis (6), and lipid composition (41) are also important for the FA mechanism.

The model directly employed and regenerated key experimental data measured at FAs. These include the size of single integrin clusters (10), the integrin activation force (6), the stretching size of single talin rods (5), the traction force vs. substrate stiffness and vinculin adhesion length vs. substrate stiffness responses of living cells (8). It was remarkable that, by simply assuming the pulling of lipid membranes at FAs, those measured data mutually supported their validity. Similar membrane analyses performed for another mechanosensitive system (42) may provide additional supports on this work, by suggesting that the mechanical response of lipid membranes is commonly important for signaling systems mediated by adhesion receptors. Overall, with an emphasis on the mechanics of lipid membranes, the presented theoretical work provides an integrated molecular mechanism for the generation of cellular traction forces and the signaling transduction at FAs.

## Supporting Materials

**Figure S1.**
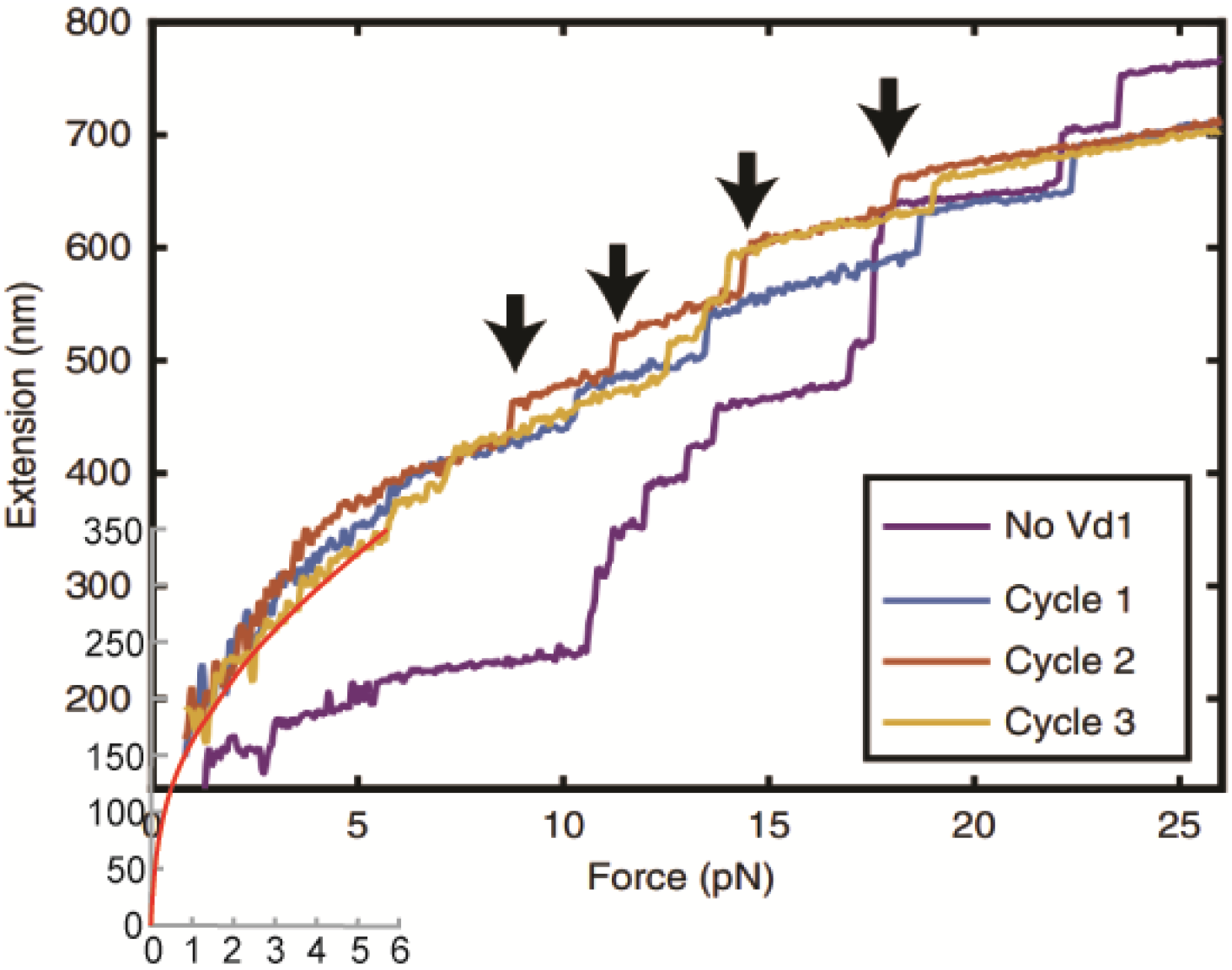
Extension vs. force response of single talins. The extension vs. force response of single full length talins was plotted by smoothly extrapolating a measured data on a larger extension regime (red). MATLAB’s smoothingspline fitting function was used. Figure for the experimental data was reprinted from (9).

**Figure S2.**
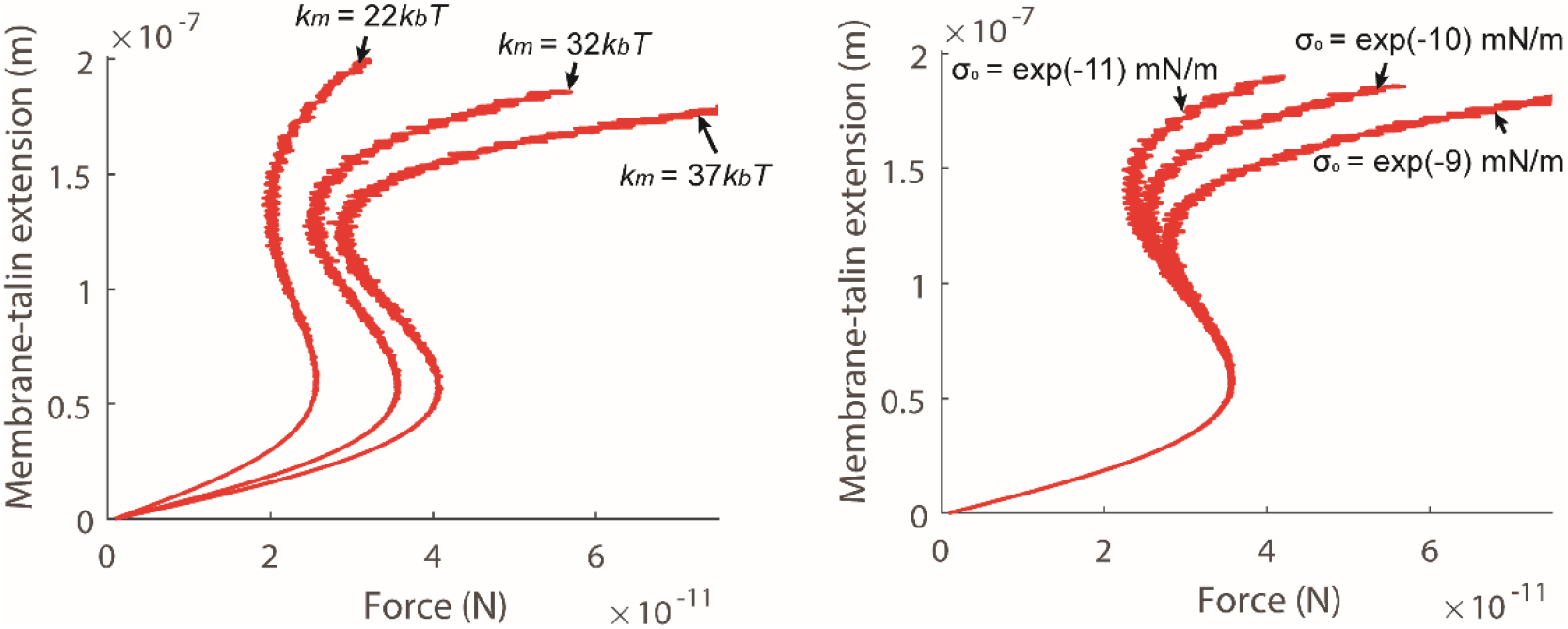
Sensitivity of *k_m_* and *σ_0_* on the extension vs. force response of the membrane-talin complex. *r_ca_* = 57 nm, *r_ct_* = 158 nm and *r_cr_* = 1750 nm were used.

**Figure S3.**
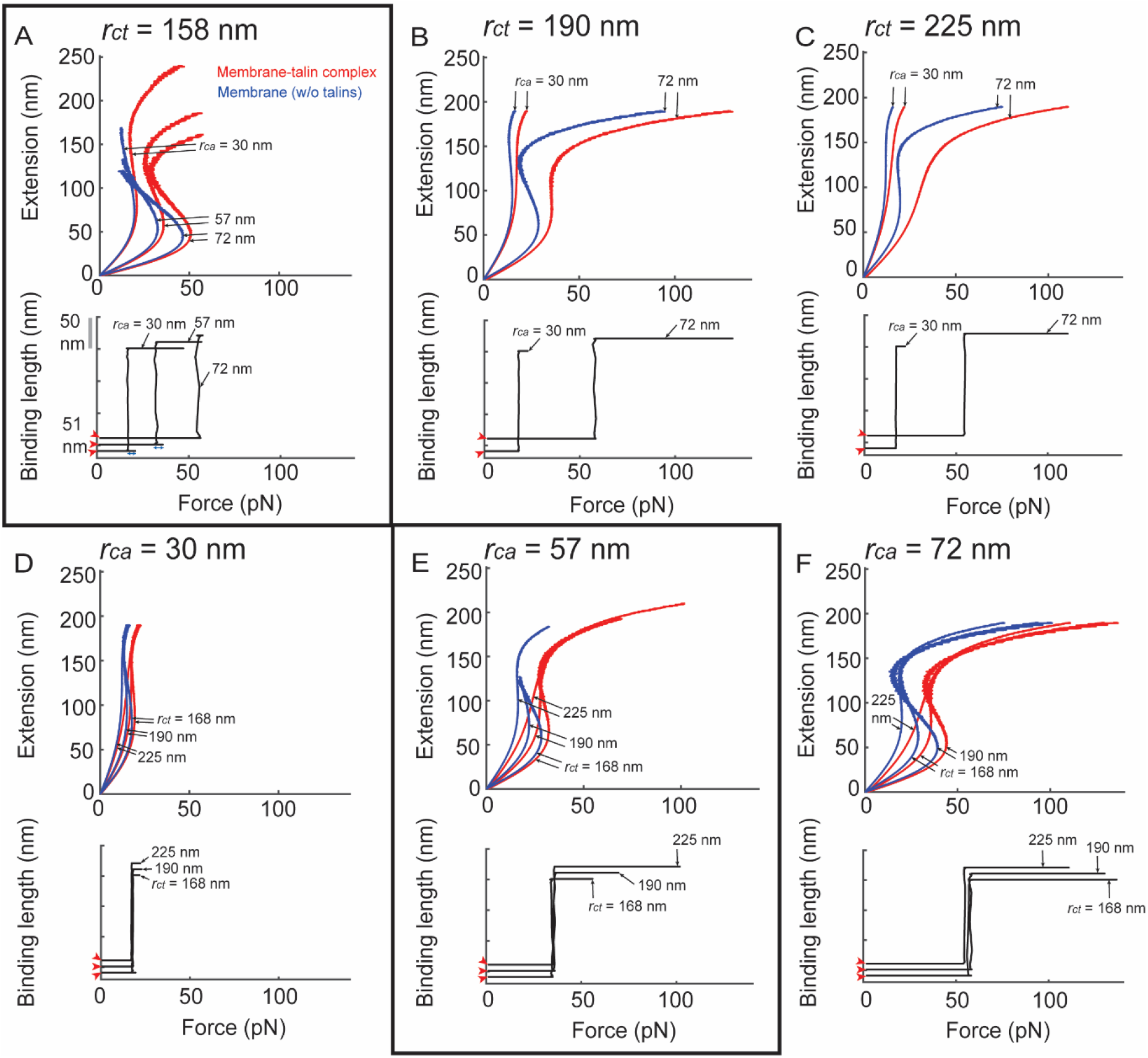
Sensitivity of *r_ca_* and *r_ct_* on the extension vs. force and vinculin binding length vs. force curves of the membrane-talin complex. (A-C) Sensitivity of *r_ca_*. *r_ct_* = 158 nm; Δ*G* = 638.4*k_b_T* and *N_talin_* ≈ 3 (*r_ca_* = 30 nm); Δ*G* = 1069*k_b_T* and *N_talin_* ≈ 9 (*r_ca_* = 57 nm); and Δ*G* = 1445.2*k_b_T* and *N_talin_* ≈ 15 (*r_ca_* = 72 nm) for (A). *r_ct_* = 190 nm; Δ*G* = 525.4*k_b_T* and *N_talin_* ≈ 3 (*r_ca_* = 30 nm); and Δ*G* = 1248.7*k_b_T* and *N_talin_* ≈ 15 (*r_ca_* = 72 nm) for (B). *r_ct_* = 225 nm; Δ*G* = 427.1*k_b_T* and *N_talin_* ≈ 3 (*r_ca_* = 30 nm); and Δ*G* = 1042.7*k_b_T* and *N_talin_* ≈ 15 (*r_ca_* = 72 nm) for (C). (D-F) Sensitivity of *r_ct_*. *r_ca_* = 30 nm; Δ*G* = 601.3*k_b_T* and *N_talin_* ≈ 3 (*r_ct_* = 168 nm); Δ*G* = 525.4*k_b_T* and *N_talin_* ≈ 3 (*r_ct_* = 190 nm); and Δ*G* = 427.1*k_b_T* and *N_talin_* ≈ 3 (*r_ct_* = 225 nm) for (D). *r_ca_* = 57 nm; Δ*G* = 1021.9*k_b_T* and *N_talin_* ≈ 9 (*r_ct_* = 168 nm); Δ*G* = 912*k_b_T* and *N_talin_* ≈ 9 (*r_ct_* = 190 nm); and Δ*G* = 759.7*k_b_T* and *N_talin_* ≈ 9 (*r_ct_* = 225 nm) for (E). *r_ca_* = 72 nm; Δ*G* = 1389.2*k_b_T* and *N_talin_* ≈ 15 (*r_ct_* = 168 nm); Δ*G* = 1248.7*k_b_T* and *N_talin_* ≈ 15 (*r_ct_* = 190 nm); and Δ*G* = 1042.7*k_b_T* and *N_talin_* ≈ 15 (*r_ct_* = 225 nm) for (F). A and E (boxed) are repeated in Fig. 2A and Fig. 2B, respectively.

**Figure S4.**
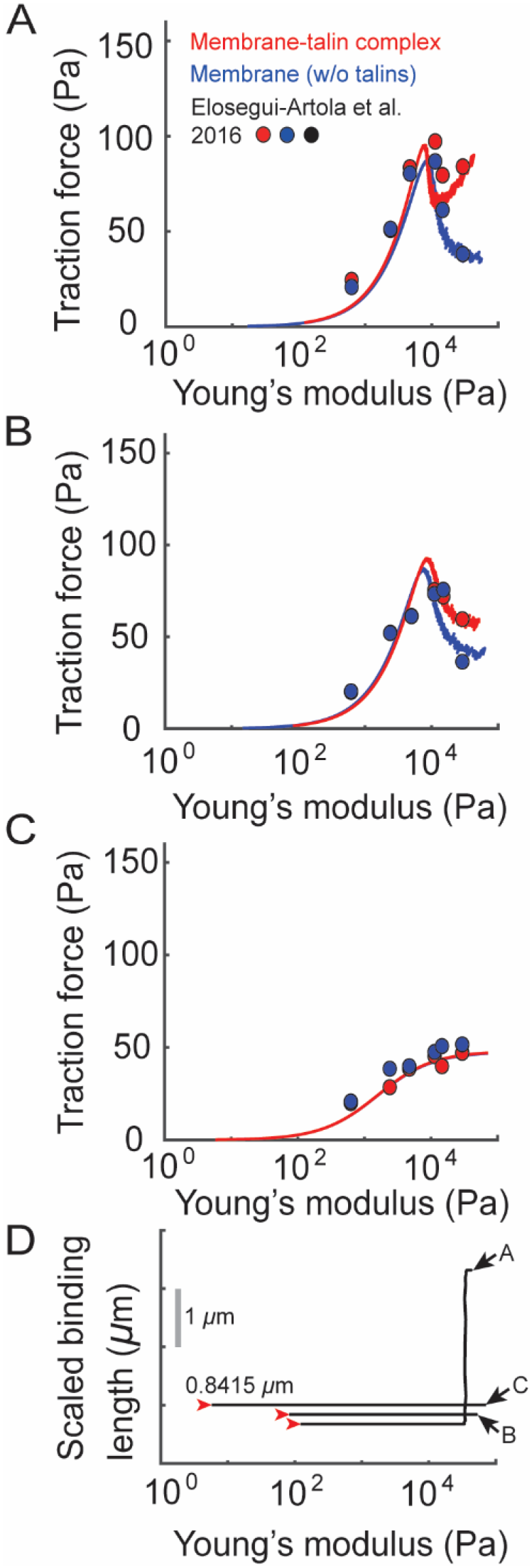
Traction force calculations by modulating *L_m_*, *C_talin_* and *r_a_*. (A-C) By using the calculation in Fig. 1C and modulating *L_m_*, *C_talin_* and *r_a_* (see Fig. S4), traction force vs. substrate stiffness responses were plotted. The membrane response (Fig. 1C, cyan) in the range from 0 nm to 132 nm and the membrane-talin response (Fig. 1C, red) in the range from 0 nm to 162.7 nm were used for (A). The range from 0 nm to 123 nm (cyan) and the range from 0 nm to 128.3 nm (red) were used for (B). The range from 0 nm to 17.41 nm (cyan) and the range from 0 nm to 17.41 nm (red) were used for (C). The calculations were matched to previous measurements obtained with the treatment of the different amounts of Blebbistatin. See Fig. S5 for the Blebbistatin concentration vs. *L_m_*/*C_g_* relation. Circles: measurements from ref. (8). Blebbistatin is known to inhibit the activity of contractile actomyosins that is closely linked to the expression level of talin proteins and the FA area (see ref. (32)).

**Figure S5.**
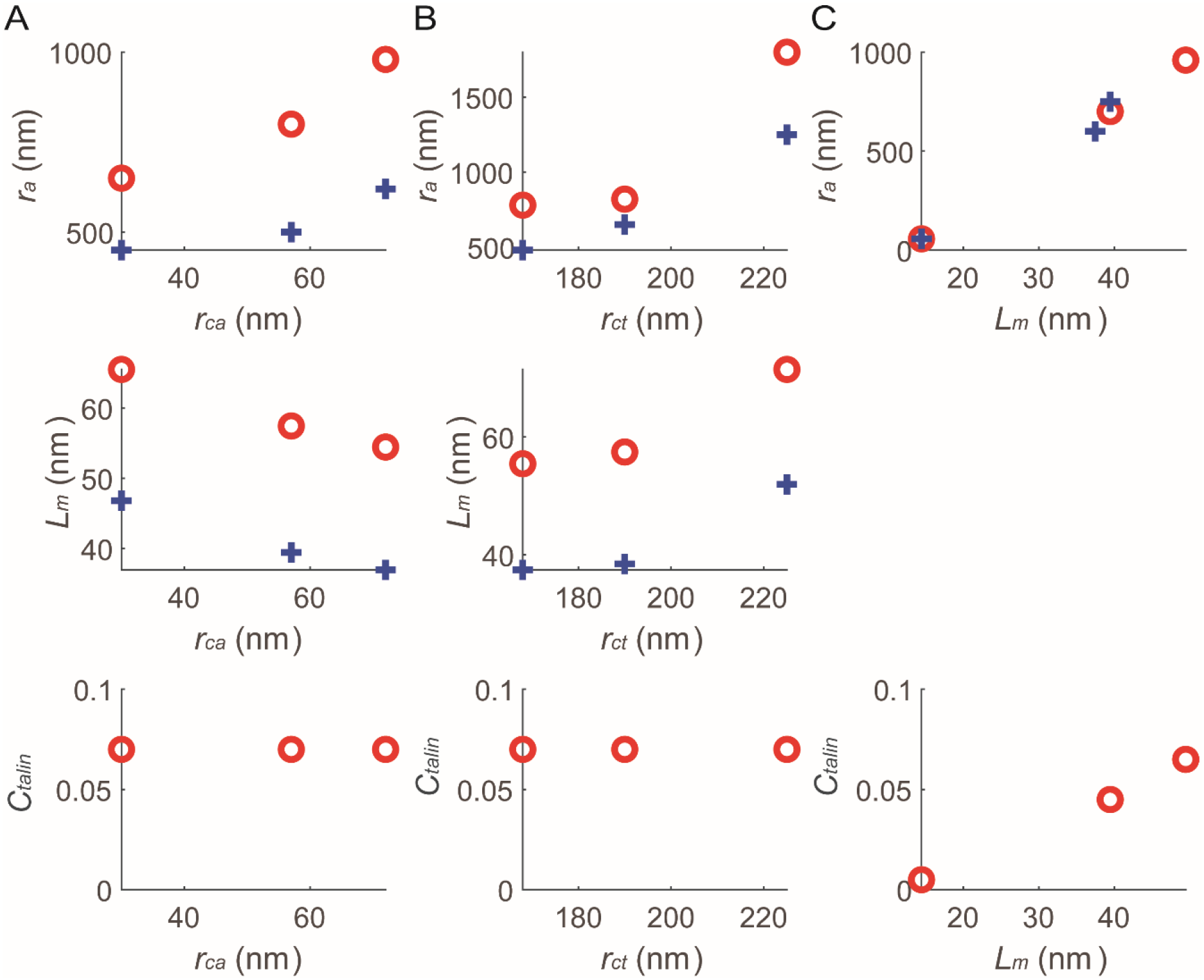
Parameter values varied in this study. (A) Values for the calculations in Fig. 3C and 3D. (B) Values for the calculations in Fig. 3E and 3F. (C) Values for the calculations in Fig. S4.

**Figure S6.**
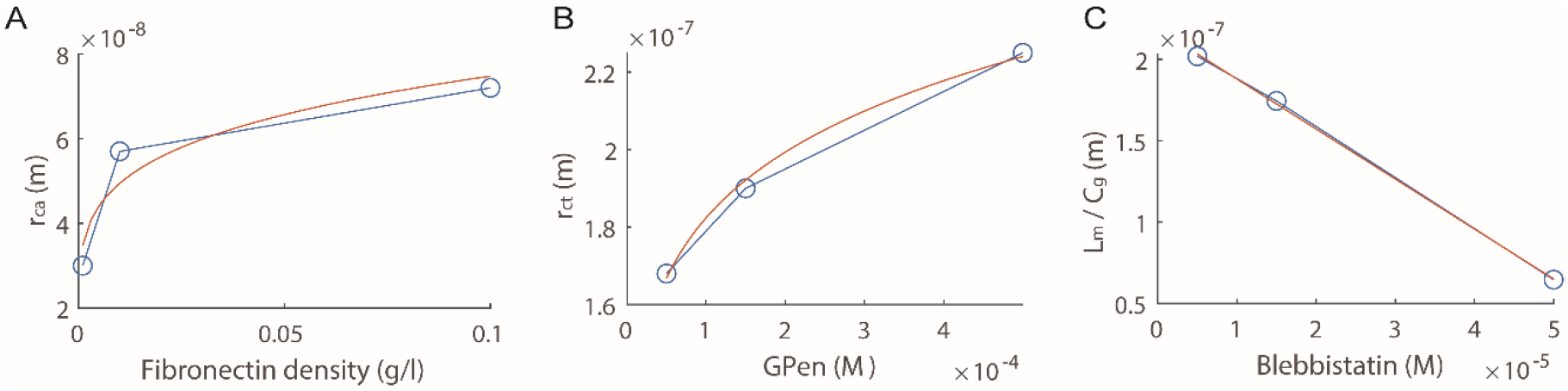
(A) The *r_ca_* vs. Fibronectin density relation. (B) The *r_ct_* vs. GPen concentration relation. (C) The *L_m_/C_g_* vs. Blebbistatin concentration relation. Best fitting analyses using MATLAB (orange) give *Y* = 1.04e-07*(*X*^0.2344^) + 1.416e-08 for (A); *Y* = 5.902e-07*(*X*^0.1194^) - 1.403e-08 for (B); *Y* = −0.003073\**X* + 2.187e-07 for (C). Experimental data were obtained from ref. (8).

